# Flagellin aerosol administration improves the efficacy of antibiotic treatment in *Actinobacillus pleuropneumoniae* infected pigs

**DOI:** 10.1101/2025.03.17.643685

**Authors:** Isabelle Fleurot, Vanaique Guillory, Céline Barc, Mickael Riou, Alain Deslis, Alexis Pléau, Thibaut Larcher, Mara Baldry, Sascha Trapp, Norbert Stockhofe, Lisette Ruuls, Conny van Solt, Jean-Claude Sirard, Ignacio Caballero

## Abstract

Swine infections with *Actinobacillus pleuropneumoniae* (*App*), the causative agent of porcine pleuropneumonia, lead to significant production losses. Surviving pigs may harbor antibiotic-resistant bacteria, highlighting the need for alternatives to reduce antimicrobial use. Stimulation of respiratory innate immunity via Toll-like receptor 5 (TLR5) agonists such as FLAMOD (a recombinant flagellin), has emerged as a promising strategy for combating bacterial lung infections. In a recent study, aerosolized administration of FLAMOD was found to elicit respiratory immune responses in pigs. This study assessed FLAMOD’s protective effects against *App*, either as a prophylactic intervention or as an adjunct to antibiotic therapy. Prophylactic administration of aerosolized FLAMOD via nebulization before *App* challenge failed to confer protection. However, in a therapeutic setting, where pigs were infected with *App* and treated with either a subtherapeutic dose of penicillin G (PenG) alone or a combination of PenG and aerosolized FLAMOD. Notably, the FLAMOD and PenG combination significantly improved bacterial clearance from the lungs. Moreover, FLAMOD-PenG treated pigs exhibited a trend toward increased weight gain and fewer lung lesions compared to those receiving PenG alone. These findings provide proof for the concept that FLAMOD enhances antibiotic efficacy, potentially lowering the required antibiotic dose and contributing to antimicrobial resistance mitigation in pigs.

## Introduction

*Actinobacillus pleuropneumoniae* (*App*), a Gram-negative coccobacillus that is facultatively anaerobic and a member of the Pasteurellaceae family, is the causative agent of porcine pleuropneumonia (1). Currently, there are 19 known *App* serovars. *Actinobacillus pleuropneumoniae* is linked to the multifactorial porcine respiratory disease complex. The severity of porcine pleuropneumonia is strongly related to *App* virulence factors including capsular polysaccharides, outer membrane proteins, exotoxins, lipopolysaccharides, permeability factors and iron-regulated proteins. The disease manifests as acute severe fibrino-hemorrhagic necrotizing lung inflammation accompanied by fibrinous pleuritis and chronic pleuritis, causing animal and production losses and presenting a significant economic burden to the pig industry worldwide (1). Although vaccination and control programs have been implemented, antibiotic therapy continues to be the standard care for controlling disease and outbreaks. However, retrospective studies have pointed to a significant increase over time in the resistance to several antibiotics (amoxicillin, amoxicillin/clavulanic acid, ampicillin, cefquinome, cotrimoxazole, penicillin G and tilmicosin) (2, 3), suggesting that pigs surviving *App* infection after antibiotic therapy may become lifelong carriers of antibiotic-resistant bacteria. In the face of escalating antibiotic resistance among *App* strains, it is imperative to explore alternative therapeutic options.

A promising option is boosting innate immune responses as a prophylactic or therapeutic measure against infectious diseases. This approach leverages the activation of Toll-like receptors (TLRs), the principal pattern recognition receptors in the innate immune system, to mobilize a broad array of host defense mechanisms that expedite microbial clearance (4). Among TLRs, TLR5 has gained particular recognition as a promising target for immunomodulatory treatments. TLR5 is located at the surface of immune and structural cells, including airway epithelial cells. It recognizes flagellin, the main component of bacterial flagella. (5). Upon stimulation, TLR5 triggers a signaling cascade that activates NF-κB, leading to an inflammatory response. This response is hallmarked by the production of pro-inflammatory cytokines and the subsequent recruitment and maturation of immune cells (6, 7). Activation of TLR5 signaling by flagellin has proven effective in providing defense against a wide range of bacterial pathogens within the respiratory tract in mice models (8, 9). This protection is linked to an early enhancement of neutrophil recruitment to the airways and the expression of antimicrobial peptides from the cathelicidin family (8, 10, 11).

There are two main intervention strategies for the use of flagellin for lung administration. The first intervention strategy involves a prophylactic delivery, where flagellin is administered prior to the bacterial challenge (e.g. with *Streptococcus pneumoniae* or *P. aeruginosa*) to stimulate host defenses as shown in mice (8, 9, 11–13). In the pig model, preventive administration of flagellin into the lungs induced a transitory cytokine response that was followed by increased numbers of inflammatory cells in the lungs. (7). Furthermore, prophylactic intervention prior to *Pseudomonas aeruginosa* infection dampened the mucosal immune response in the airways, both *in vitro* and *in vivo*, thereby reducing the severity of lung lesions in infected pigs (7). The second strategy focuses on the adjunct effect of flagellin administered with antibiotics post-infection. Previous studies showed that combination of antibiotic treatment with flagellin in a therapeutic setting enhances the antibacterial efficacy of antibiotics against antibiotic-sensitive and -resistant *Streptococcus pneumo*niae (14–17) and *Klebsiella pneumoniae* (18) in mice models of pneumonia. This positive effect of the combined therapy highlights flagellin’s potential in combating the rise of antibiotic resistance.

The aim of the present study was to evaluate whether flagellin, administered either prior to an *App* challenge or post-infection in combination with an antibiotic, confers protection against *App* infection. The findings obtained provide proof for the concept that flagellin administration can enhance antibiotic efficacy in an experimental model of *App* infection.

## Materials and Methods

### Animals

Animal experiments to test the prophylactic effect of flagellin against *App* were conducted in accordance with the Dutch animal experimental and ethical requirements and the project license application was approved by the Dutch Central Authority for Scientific Procedures on Animals (AVD4010020187205) and the experiment plan was approved by the institute’s Animal Welfare Body (Permit number: 2018.D-0042). Twenty-four male six-week-old pigs (TOPIGS 20 breed) were housed in biosafety level 2 (BSL-2) containment housing throughout the whole procedure at Wageningen Bioveterinary Research in Lelystad, Netherlands. The pigs were housed in a HEPA-filtered facility and distributed among three pens, each providing approximately 1 m² of floor space per animal. Pens contained some straw as floor bedding and were enriched with toys as playing material. Drinking water was freely available by drinking nipples. Commercial feed for pigs in nursery units was provided ad libitum. The experiment started after one week of acclimatization.

Experimental *in vivo* infections to test the therapeutic effect of a combined therapy of flagellin and antibiotics post-*App* infection were performed at the INRAE PFIE animal experimental platform (UE-1277 PFIE, INRAE Centre de Recherche Val de Loire, France, https://doi.org/10.15454/1.5535888072272498e12<u>; member of the national infrastructure</u>) and approved by the local Ethics Committee in animal experimentation Val de Loire, CEEA-019 (reference number APAFIS#30918-2021040618206954 v3). A total of 32 healthy 1-month-old Large White pigs were used in these experiments. Pigs were kept in BSL-2 housing (9 m2/pig) that was cleaned daily throughout the experimental procedure. Animals had access to a standard grain-based diet (Sanders) and water ad libitum. Social and material (balls) enrichment was provided to maintain pig welfare. The physical condition and animal welfare were determined as described in (19).

### Production of recombinant flagellin

The recombinant flagellin FliC_Δ174-400_ used in this study, *i.e.,* FLAMOD derives from *Salmonella enterica* serovar Typhimurium FliC. The recombinant flagellin FliC_Δ174-400_ lacks the antigenic domain from the flagellin. The molecule has a lower intrinsic antigenicity, compared to native flagellin, and it has been demonstrated to be safe even after several repeated flagellin administrations (14, 20). It was produced in inclusion bodies in *Escherichia coli* by the Vaccine Development department at Statens Serum Institut, Denmark. This flagellin was purified by filtration and chromatography and resuspended in the diluent buffer 10 mM phosphate, 145 mM NaCl, polysorbate 80 0.02 % (w/v) pH 6.5. The buffer was defined to maintain the integrity and pharmacological activity of flagellin during mesh-nebulisation by the Aerogen Solo (Aerogen) (15). The immunostimulatory activity was tested using the HEK-Dual™ hTLR5 cells assay (Invivogen). The low endotoxin content in the protein preparation was assessed with a *Limulus* assay (Pyrochrome, kinetic LAL assay from Associates of Cape Cod Inc.).

### Prophylactic administration of flagellin prior to *Actinobacillus pneumoniae* challenge

Pigs were randomized in an unbiased manner and placed in 3 different groups (n=8/group). The first group consisted of animals infected with *App*. The second group received FLAMOD by nebulization and 24h later was infected with *App*. The third group was infected with *App* and treated with a therapeutic dose of intramuscular penicillin G (300.000 IU/mlDepocillin®, MSD Animal Health) once per day, starting directly after infection.

For aerosol administration of FLAMOD, pigs were anesthetized by intramuscular injection of Tiletamine-Zolazepam (Zoletil®250, 50 mg/ml, Virbac) and Xylazinehydrochloride (Sedaxylan® 20mg/ml, Dechra, NL) at 0.1 ml/kg and laid into a half pipe in sternal position. Aerosol delivery was performed using an Aerogen mesh nebulizer system. Animals that did not receive FLAMOD were also anesthetized to avoid confounding effects of the anesthesia. The aerosol was generated from 2 mL of FLAMOD solution at a 1.5 mg/ml concentration. Twenty-four hours later, animals were challenged with 2 ml of an *App* serotype 2 suspension at a concentration of 1 x 10^6^ CFU/ml using a MAD Nasal™ Intranasal Mucosal Atomization Device (Teleflex®) and sacrificed 48h p.i. by exsanguination for sample collection. The animals were monitored daily following infection to assess disease progression and discomfort. Humane endpoints were predefined to avoid unnecessary animal suffering that required euthanasia. Body weight and temperature were monitored twice daily.

### Therapeutic administration of FLAMOD post *Actinobacillus pleuropneumoniae* challenge

To assess the therapeutic efficacy of FLAMOD against *App* infection, we first aimed to establish a severe *App* pig infection model. Thirteen 1-month-old Large White pigs were infected with 2 ml of an *App* serotype 2 suspension at a concentration of 5 x 10^6^ CFU/ml (n=9) or sham-infected (n=4) using a MAD Nasal™ Intranasal Mucosal Atomization Device (Teleflex®). In the next set of experiments, we aimed to evaluate whether flagellin treatment could improve the efficacy of antibiotic treatment in *App* infected pigs. Pigs were infected as described above, nebulized with 1.3 mg of FLAMOD (n=10) or the vehicle (n=9) using an Aerogen mesh nebulizer system at 3h p.i. and treated with a sub-therapeutical antibiotic dose (4000 UI/kg Depocilline®; active compound penicilling G [PenG]) at 3, 24 and 48h p.i. The animals were monitored for a period of 3 days following infection to assess disease progression. Animals reaching predefined humane endpoints, as determined by established clinical or behavioral criteria indicative of severe distress or suffering, were euthanized prior to the trial’s end to minimize unnecessary suffering. Body weight and temperature were monitored at 0, 3, 6, 24, 48 and 72h p.i.

### Pathology analysis of the lungs

Animals were anaesthetised with a mixture of zoletil/xylazine and euthanized by exsanguination. The lungs were carefully removed for examination of the upper and lower respiratory tract. Macroscopic lesions were evaluated using a previously published scoring system where each of the seven lung lobes were given a score (0 to 5) based on the number and size of lesions with a maximum total score of 35 (21). For histological evaluations, sections (5 µm) of 4% formalin-fixed and paraffin-embedded samples were stained with hemalum-eosin saffron (HES). Histological findings were evaluated blindly by a certified pathologist. Briefly, 9 lung elementary lesions were identified (lesion extension, emphysema, alveolar wall infiltration, alveolar epithelium damage, alveolar haemorrhages, alveolar exudate, epithelium damage, bronchial occlusion and vascular lesions). Scores were given from 0 (no findings) to 3 (extended manifestation).

### Bacteriological analysis

Bacterial loads were analysed from lung samples collected at the same locations (i.e. cranial left and cranial right lung lobes) in selective Chocolate agar PolyViteX media (Biomérieux). Blood cells were counted at 0, 6 and 72h p.i. using a MS9-5 Hematology Counter® (digital automatic hematology analyzer, Melet Schloesing Laboratories, France).

### ELISA

To explore changes in the inflammatory status of the infected animals, a Luminex immunoassay was performed on serum samples from infected pigs treated with either a combination of antibiotic plus flagellin or solely the antibiotic at 0, 6 and 7h p.i. The Porcine Cytokine/Chemokine Magnetic Bead Panel kit (Merck Millipore) was used to measure the following cytokines: GM-CSF, IFN-γ, IL-1α, IL-1β, IL-1ra, IL-2, IL-4, IL-6, IL-8, IL-10, IL-12, IL-18, and TNF-α. Measurements were made using a Luminex LX200 reader (Merck Millipore) following the manufacturer’s recommendations. In addition, the acute phase C-reactive protein (CRP) was measured in serum using a Pig CRP ELISA Kit (Abcam) in accordance with the manufacturer’s instructions.

### RNA sequencing

Total RNA was extracted from the cranial left lung lobe using the Nucleospin® RNA Set for NucleoZol kit (Macherey-Nagel, Düren, Germany) according to the manufacturer’s protocol. RNA integrity was checked with the Bioanalyzer RNA 6000 Nano kit (Agilent Technologies). The NEBNext® Ultra™ II RNA Library Prep Kit for Illumina® was used to prepare the RNA library. Sequencing was performed on a Illumina NovaSeq 6000 platform (Illumina) with 2[×[150bp paired-end chemistry. Raw reads were trimmed with Trim Galore. Read quality assessment was performed using FastQC and FastQ Screen tools to mine for potential contamination. Reads were then mapped and quantified using STAR_2.6.1c and RSEM (v1.3.1) using the function rsem-calculate-expression (parameters: –star –sort-bam-by-coordinate) on a reference file containing the genomes of Sus scrofa (version 11.2). The produced count matrix was then used for all subsequent statistical analysis within the statistical package R version 4.3.2. Differential gene expression was calculated using the DESeq function from the Bioconductor package DESeq2 (v1.42.1) (22). For each gene, three negative binomial generalized linear model were fit. The first contained as parameter the comparison between the combined antibiotic (PenG) and FLAMOD therapy vs standalone antibiotic. The second compared differential gene expression between animals with high lesion score (score higher or equal than 6) and low lesion score (score lower than 6). The third model compared animals with high bacterial count (log10 CFU higher than 4) vs animals with low bacterial count (log10 CFU equal or lower than 4). Additional parameters were added to both models to account for bias due to the date of the experiment and the sex of the animals. We considered differentially expressed genes those with p-adjusted value <0.05 and log2 fold change >1. The clusterProfiler package (23) was used to perform enrichment analysis for GO terms and KEGG pathways using a p value cutoff of 0.05 and the Benjamini-Hochberg (BH) method to adjust for multiple comparisons. Sequencing data have been deposited in the Gene expression omnibus repository with the accession number GSE253855.

### Statistical analysis

All analyses were performed using the statistical package R version 4.2.3 and the packages ggplot2, ggpubr, survival, survminer and lme4. Time-series figures are presented as mean ± SEM. A pairwise Wilcoxon test with Holm correction for multiple comparisons was used to determine significant differences between pigs pre-treated with flagellin or not and infected with *App*. A mixed linear model was fit to evaluate differences between the treatment group, time and the interaction in the time series. The pig ID was introduced as random variable. A parametric t-test was used to compare significant differences when comparing two groups. A p-value <0.05 was considered statistically significant.

## Results

### Prophylactic administration of FLAMOD did not protect against *Actinobacillus pleuropneumoniae* infection

We have previously observed that intrapulmonary inoculation of FLAMOD induces a transient cytokine release, leading to the recruitment of immune cells into the lungs (7). Similar observations were conducted upon administration of FLAMOD into lungs using nebulization. We hypothesized that early recruitment of immune cells would enhance the host defense against *App* in the porcine model. To test this hypothesis, Pigs were treated with FLAMOD and challenged via the respiratory route with a pathogenic infectious dose of *App* at 24 h. Control groups included pigs that did not receive FLAMOD treatment and were infected with *App* and pigs infected with *App* that received PenG treatment at therapeutic doses. Our data showed that aerosolized administration of FLAMOD into the lungs 24h prior to *App* challenge did not protect animals from infection. Pigs pre-treated with FLAMOD prior to *App* infection had increased temperature compared to animals receiving PenG and similar body weight gain as non-treated infected pigs and non-treated infected pigs receiving PenG therapy (Figures 1A-B). Three out of the 8 (37.5%) non-treated infected pigs presented lung lesions, whereas 5 out of 8 pigs pre-treated with FLAMOD (62.5%) developed *App* lesions. None of the PenG-treated pigs displayed any signs of lesions in the lungs. In addition, the percentage of lung area affected by lesions was significantly increased in the FLAMOD-pre-treated animals compared to those treated with PenG (Figure 1C). *App* bacteria were isolated from 7 lungs in the non-treated group and from 8 lungs in the FLAMOD treated group. Finally, no significant differences were observed in the percentage of macrophages and neutrophils among the three groups (Supplementary figure 1). These results suggest that FLAMOD administration prior to *App* infection does not enhance protection against *App*.

**Figure 1.**
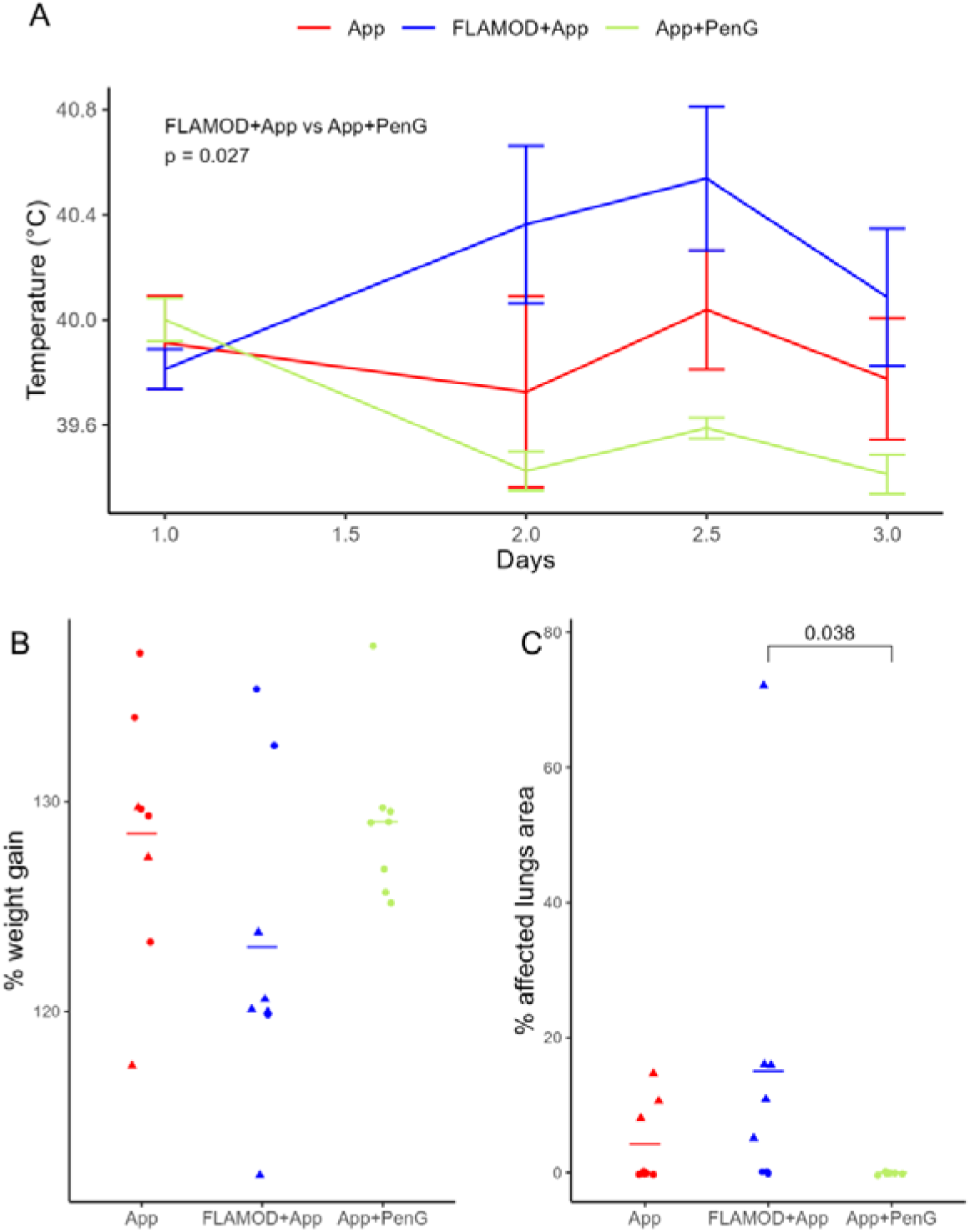
FLAMOD pre-treatment does not protect against *App* infection. Pigs were pre-treated (FLAMOD+*App*) or not (*App*) with 2 mL of FLAMOD solution at a 1.5 mg/ml concentration for 24h before being challenged with of 2 x 10^6^ CFU of an *App* serotype 2 and then sacrificed 48h p.i. A control group where the pigs were infected as before and treated with a therapeutic dose of penicillin G (*App*+PenG) was included. (A) Body temperature. (B) Percentage of weight gain during the experiment. (C) Percentage of affected lungs area. A pairwise Wilcoxon test with Holm correction for multiple comparisons was used to evaluate significant differences between the FLAMOD+*App*, *App* and *App*+PenG grouprs. Triangle shapes indicate animals that presented lung lesions. Data are representative of 24 animals.

### Combined administration of FLAMOD and penicillin G improves the treatment of *Actinobacillus pleuropneumoniae* infection

To test the effect of a combined FLAMOD-PenG treatment against *App* infection, we first aimed to establish a more severe model of *App* infection, as only 37.5% to 62.5% of pigs showed pathological lesions in our previous prophylactic setting experiment. To this end, the dose of the *App* inoculum was increased about 5-fold. One-month-old Large White pigs were infected with a single intranasal administration of 2 ml of 5 x 10^6^ CFU/ml of *App* or the vehicle. Pigs were monitored for 3 days before being sacrificed for sample collection. We observed that infected animals developed respiratory distress and 5 out of 9 animals needed to be sacrificed before the end of the trial (Figure 2A). Infected animals developed lesions characteristic of acute *App* infection, including hemorrhagic pneumonia, multifocal necrotic foci and fibrinous exudation (Figures 2B-C). At the histological level we could observe the presence of alveolar damage, vascular damage, fibrin accumulation and an extensive influx of inflammatory cells (Figure 2D-E). In conclusion, our findings were consistent with a severe to fatal infection leading to the death of 55% of the animals.

**Figure 2.**
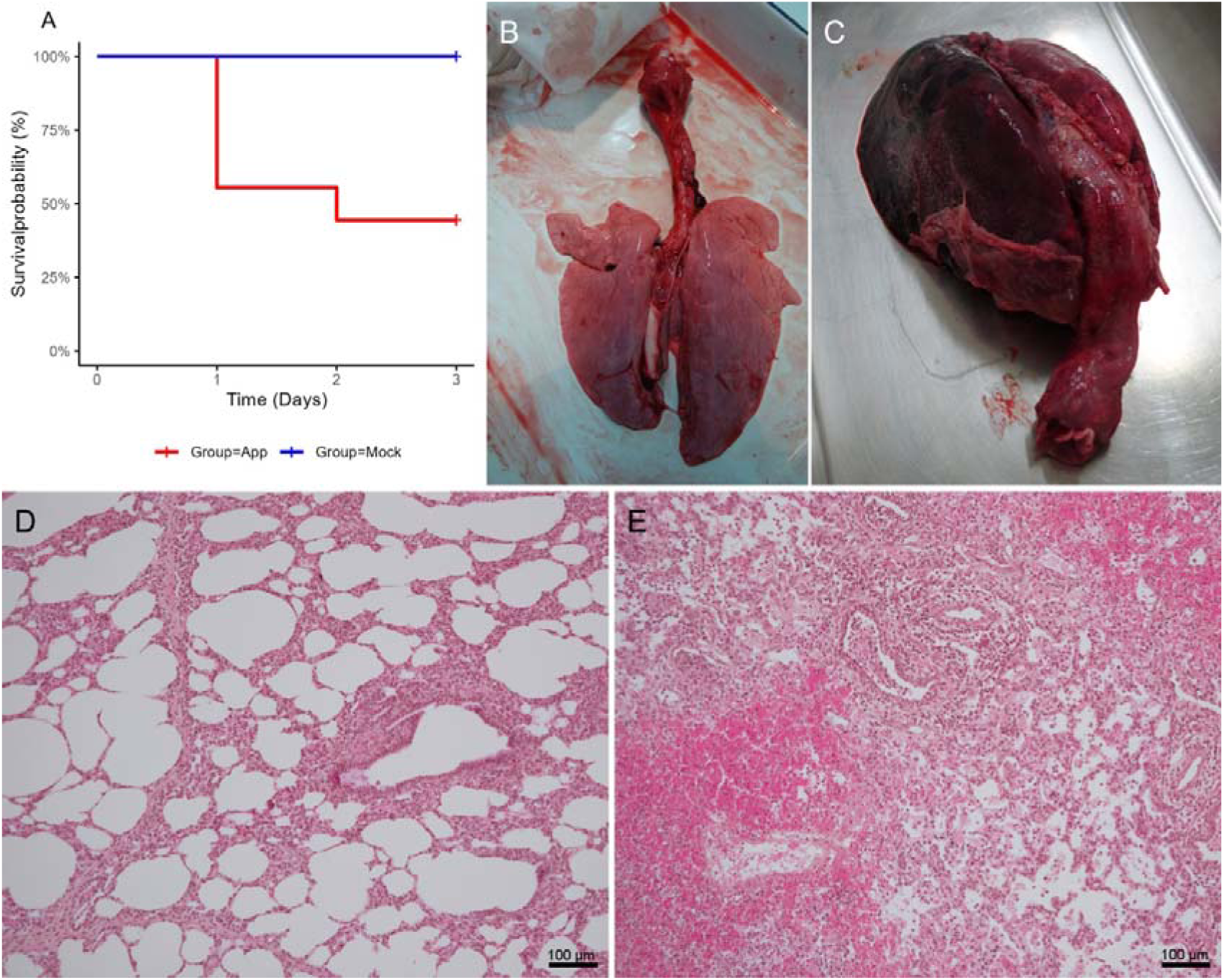
*Actinobacillus pleuropneumoniae* challenge leads to severe pneumonia. One-month old pigs were intranasally inoculated with either 1 x 10^7^ CFU *App* serotype 2 or PBS. (A) Animal survival recorded over a period of 3 days post-infection. (B) Representative image of the lungs from sham-infected pigs (Mock). (C) Representative image of the lungs from pigs infected with *App.* Lungs presented hemorrhagic lesions with necrotic tissue and fibrinous exudation. (D-E) Lung sections (left cranial lobe; 10x) were stained 3 days post-infection with hemalum-eosin saffron. Images are shown of (D) sham-infected and (E) *App*-infected pigs. Data are representative of 12 animals.

Once we established an experimental infection model that leads to severe disease, we evaluated whether the combined therapy of PenG and FLAMOD (PenG-FLAMOD) protects against *App*. Pigs were infected with *App* as described above and then treated with PenG or a single administration of nebulized FLAMOD as adjunct of PenG treatment. All animals treated with PenG or the combined therapy of PenG-FLAMOD survived the *App* challenge infection. We observed a slight drop in body weight at 24h p.i., however, the animals recovered and had a significant gain in weight at the end of the experimental period (Figure 3A). Interestingly, the animals treated with the PenG-FLAMOD combination had a better recovery and showed a tendency towards greater weight gain compared to animals treated only with PenG (Figure 3A). There were no significant differences in body temperature between PenG-FLAMOD treated and PenG-treated pigs (Figure 3B). We observed an increase in circulating monocytes in the pigs that were administered the combination treatment (PenG-FLAMOD) at 6h p.i. (which corresponds to 3h post-FLAMOD administration), compared to their PenG-treated counterparts (Figure 3C). No differences were observed in any of the other clinical parameters assessed.

**Figure 3.**
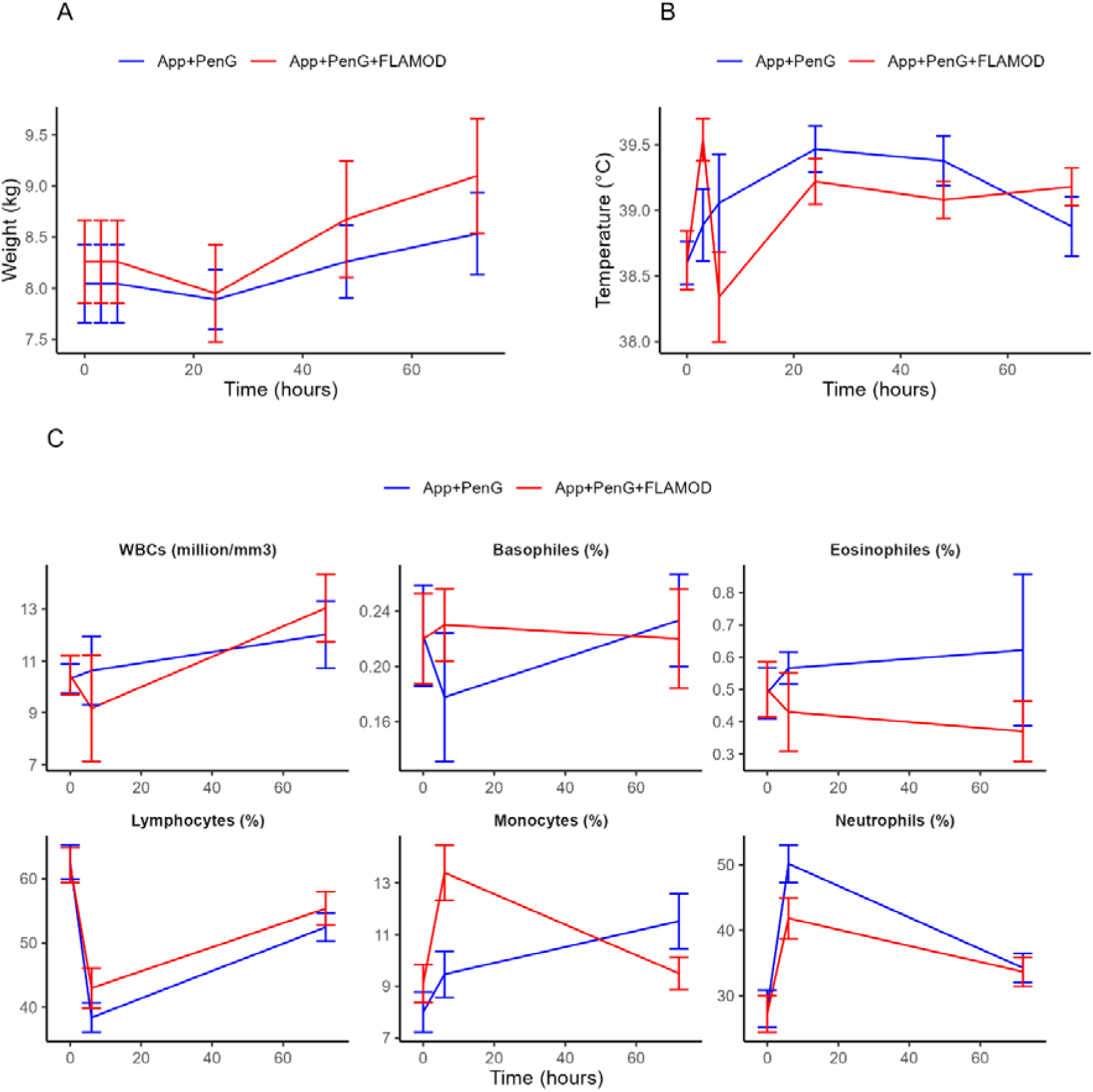
Clinical parameters and white blood cell count in *App* infected pigs are similar between animals treated with antibiotic (*App*+PenG) or a combination of antibiotic and FLAMOD (*App*+PenG+FLAMOD). Pigs were intranasally inoculated with 1 x 10^7^ CFU *App* serotype 2 and then treated or not with 1.3 mg of FLAMOD 3h p.i. In addition, pigs were treated with a daily intramuscular injection of 4mg/kg penicillin G. Evolution of the (A) animal’s weight and (B) body temperature throughout the experimental period. (C) Total white blood cell (WBCs) in peripheral blood. A mixed linear model was fitted to evaluate differences between the treatment group, time and the interaction in the time series. The pig ID was introduced as random variable. Data are representative of 9-10 animals per experimental group.

The severity of macroscopic lung lesions varied significantly, ranging from no detectable lesions to readily visible subacute or hyper-acute manifestations. In the more severe cases, these lesions were characterized by pulmonary consolidation, along with the occurrence of numerous necrotic areas, hemorrhages, and the accumulation of fibrous exudation (Figure 4A-B). Histological evaluation showed the presence of lesions and inflammatory cells in both PenG-treated and PenG-FLAMOD-treated pigs. The alveolar lumen was filled with fibrin and immune cells with an extensively thickened alveolar wall (Figure 4C-D). Nevertheless, no differences were observed at the histological level between the two treatment groups. Despite the great individual variability between animals in the severity of the pathological lesions, we observed a tendency towards a decreased lesion score at the macroscopic level in animals that received the PenG-FLAMOD combination treatment (Figure 4E). Samples were collected from the same area of the left and right cranial lobe 3 days p.i. in all animals for bacteriological analysis. Importantly, the number of bacteria in the lungs dropped significantly when pigs were treated with the PenG-FLAMOD combination therapy compared to those treated solely with the antibiotic (median 995.04 CFU vs 124743.6 CFU in PenG-FLAMOD treated vs PenG-treated, respectively; Figure 4F). Altogether, these data suggest that FLAMOD improves the general health status and the efficacy of antibiotic therapy in *App* infected pigs.

**Figure 4.**
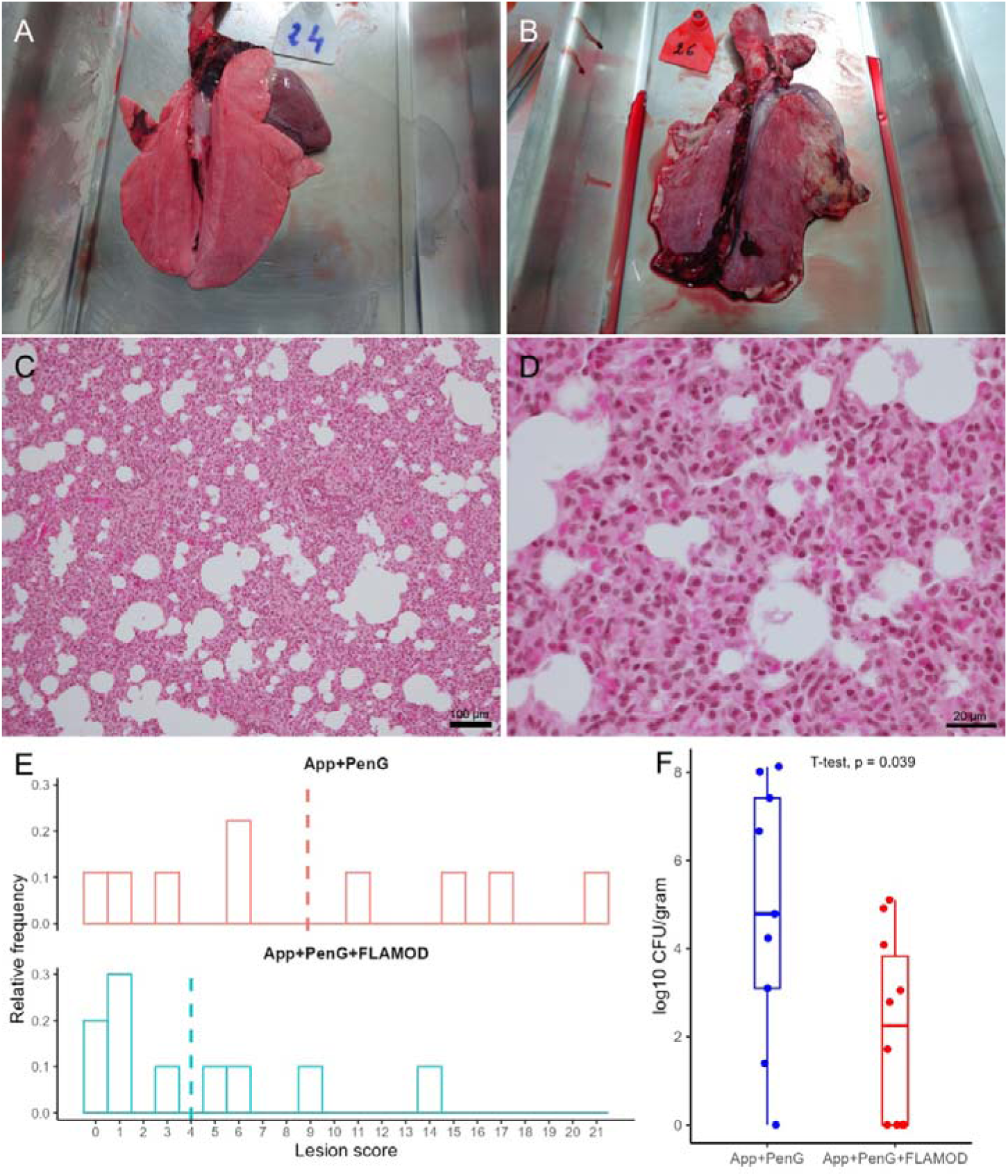
FLAMOD improves the antibiotic treatment effect in the *App* challenge infection. Pigs were inoculated intranasally with 1 x 10^7^ CFU *App* serotype 2 and treated with a daily dose of penicillin G (*App*+PenG) or a combination of a daily dose of penicillin G and a single administration of 1.3 mg of FLAMOD (*App*+PenG+FLAMOD). (A-B) Examples of pig lungs presenting low and high score macroscopic lesions. (C-D) Histological images stained with hemalum-eosin saffron. (C) We observe the presence of inflammatory cells or red blood cells secondary to vascular damages, along with thickened alveolar wall due to inflammatory cell infiltration (left cranial lobe; 10x). (D) Higher magnification (left cranial lobe; 40x) showing alveoli with fibrin deposits and enlarged alveolar walls with inflammatory cells. (E) Histogram showing the distribution of the lesion score in *App*+PenG and *App*+PenG+FLAMOD pigs evaluated at day 3 p.i. using the score system developed by Hannan et al. (21). Dashed lines indicate the mean value of the lesion scores. The y-axis represents the relative frequency, calculated as the number of events in each bin divided by the total number of events. (F) Determination of bacterial counts in the lungs at day 3 p.i. A parametric t-test was used to compare differences between *App*+PenG and *App*+PenG+FLAMOD groups. Data are representative of 9-10 animals per experimental group.

### Nebulization of FLAMOD modulates the production of circulating cytokines during infection

Previous studies have demonstrated that flagellin exerts an immunostimulatory effect in pig lungs when administered by the respiratory route. We evaluated the production of circulating cytokines and CRP in the pig serum at 0, 6 and 72h p.i. (Figure 5). The combination therapy of PenG and FLAMOD significantly increased and maintained elevated levels of IFN-γ, IL-1α, IL-4, IL-10, IL-12, and IL-18 during the entire 72h of experimentation. In contrast, PenG treatment alone induced only an early transient rise in cytokines followed by a subsequent decline. The PenG-FLAMOD treatment was also associated with a significant transitory increase in IL-8 at 6h p.i. These data suggested that respiratory delivery of FLAMOD also impact systemic immune responses. Finally, CRP was elevated in infected pigs treated solely with PenG at 72h post-infection. CRP levels were significantly lower in animals that were treated with the PenG-FLAMOD combination therapy, suggesting a rapid resolution of the inflammatory process (Figure 5 and supplementary figure 2).

**Figure 5.**
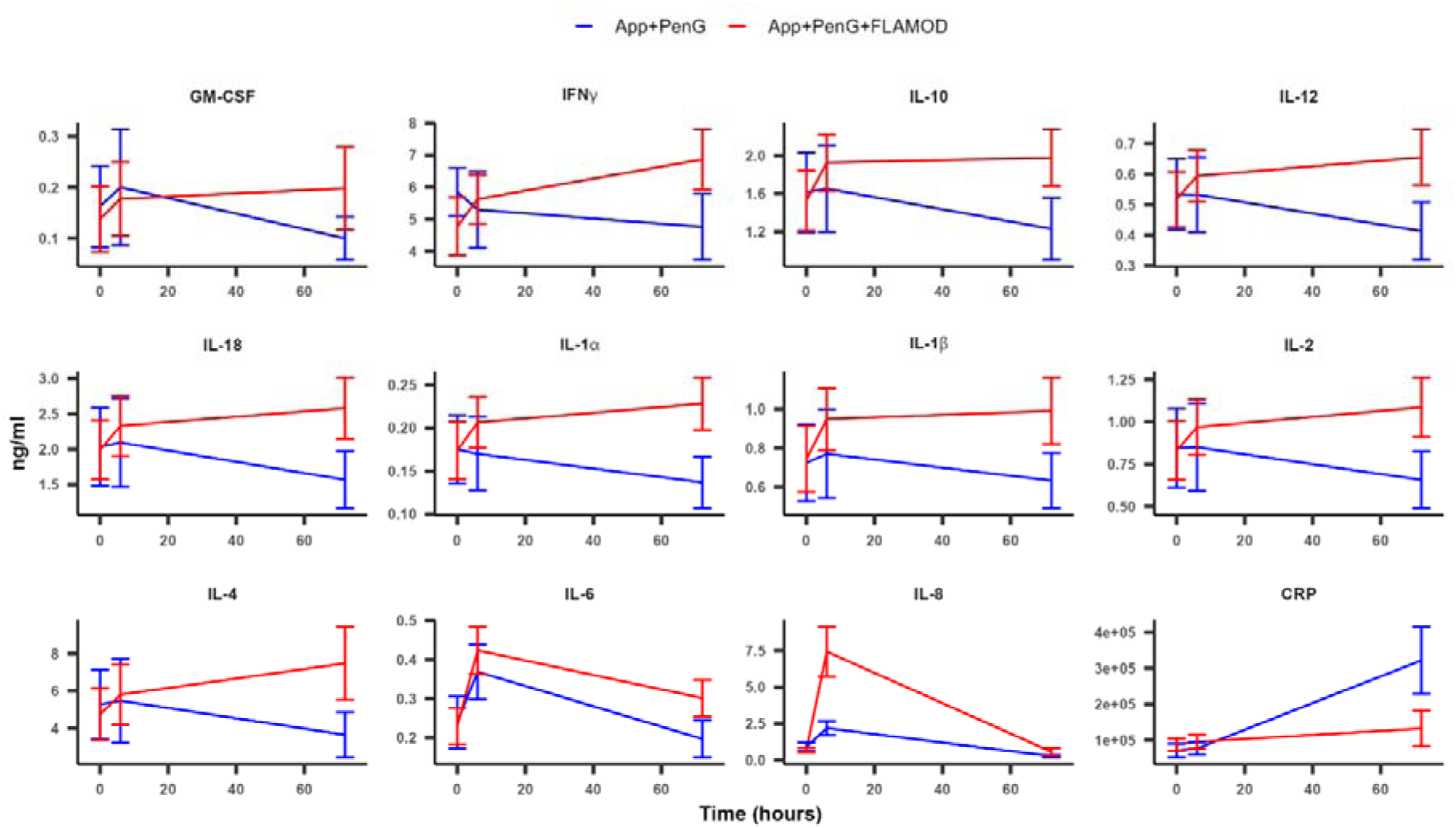
Effect of the combination therapy of FLAMOD and penicillin G on the pig circulating cytokines during *App* infection. One-month old pigs were infected with 1 x 10^7^ CFU *App* serotype 2 and treated daily with 4 mg/kg of penicilling G (*App*+PenG; n=9) or with one dose of 1.3 mg of FLAMOD in addition to the antibiotic therapy (*App*+PenG+FLAMOD; n=10). (A) Measurement of GM-CSF, IFN-γ, IL-1α, IL-1β, IL-2, IL-4, IL-6, IL-8, IL-10, IL-12, IL-18 and CRP production in the serum at 0, 6 and 72h p.i. A mixed-effects linear model was applied to analyze statistical differences among the treatment groups, temporal factors, and the interaction effect in the time series data. Pig identification was included as a random effect in the model.

### FLAMOD’s effect in the lungs is transitory while transcriptional changes are driven by the severity of the lung infection

To determine alterations in lung immunity, we performed RNA sequencing (RNA-seq) analysis on the pig lungs post-mortem (3 days p.i.). There was only 1 gene (*ANO1*) that was upregulated (padj<0.05) in lung tissue between the PenG-FLAMOD combination therapy versus standalone PenG (supplementary figure 3). To get further insights, two other comparisons were performed. We identified 33 differentially expressed genes (DEGs) when we compared animals with high bacterial loads (log10 CFU > 4) vs low bacterial loads (log10 CFU ≤ than 4). The majority of DEGs were upregulated in the group with high bacterial loads and related to immunity, such as, *IL-1*β*, ACKR1, CMTM2, CHIT1, ADAMDEC1, TIMP1, BDKRB1* (Figure 6A). Gene set enrichment analysis (GSEA) showed an activation of pathways involved in the inflammatory response and response to molecules of bacterial origin (Figure 6B). Moreover, DEG analysis between animals presenting severe lesions (score ≥ 6) vs low lesions (score < 6) revealed 587 DEGs (Figure 6C). The most upregulated genes encode several cytokines and chemokines such as, *IL-1*β, *IL-18*, *CXCL8*, *CCL8*, *CCL2*, *CCL20* and *PTN;* while downregulated genes were associated with protection against inflammation such as *SLPI*. In addition, the pathways inflammatory response and defense response were enriched in the GSEA (Figure 6D). Differential expressed genes are shown in supplementary table 1. To verify the results obtained by RNA-seq, and evaluate whether there were differences in gene expression between the cranial and caudal lobes, we performed quantitative RT-qPCR in a subset of differentially expressed genes. There were significant differences in most of the genes evaluated between animals with high lesions compared to low lesions in samples collected from the cranial lobe (supplementary Figure 4). Overall, in contrast to the early response to FLAMOD in the pig respiratory tract, there is no significant impact on gene expression at 3 days post-treatment. However, most lung DEGs are associated to the severity of inflammation and infection.

**Figure 6.**
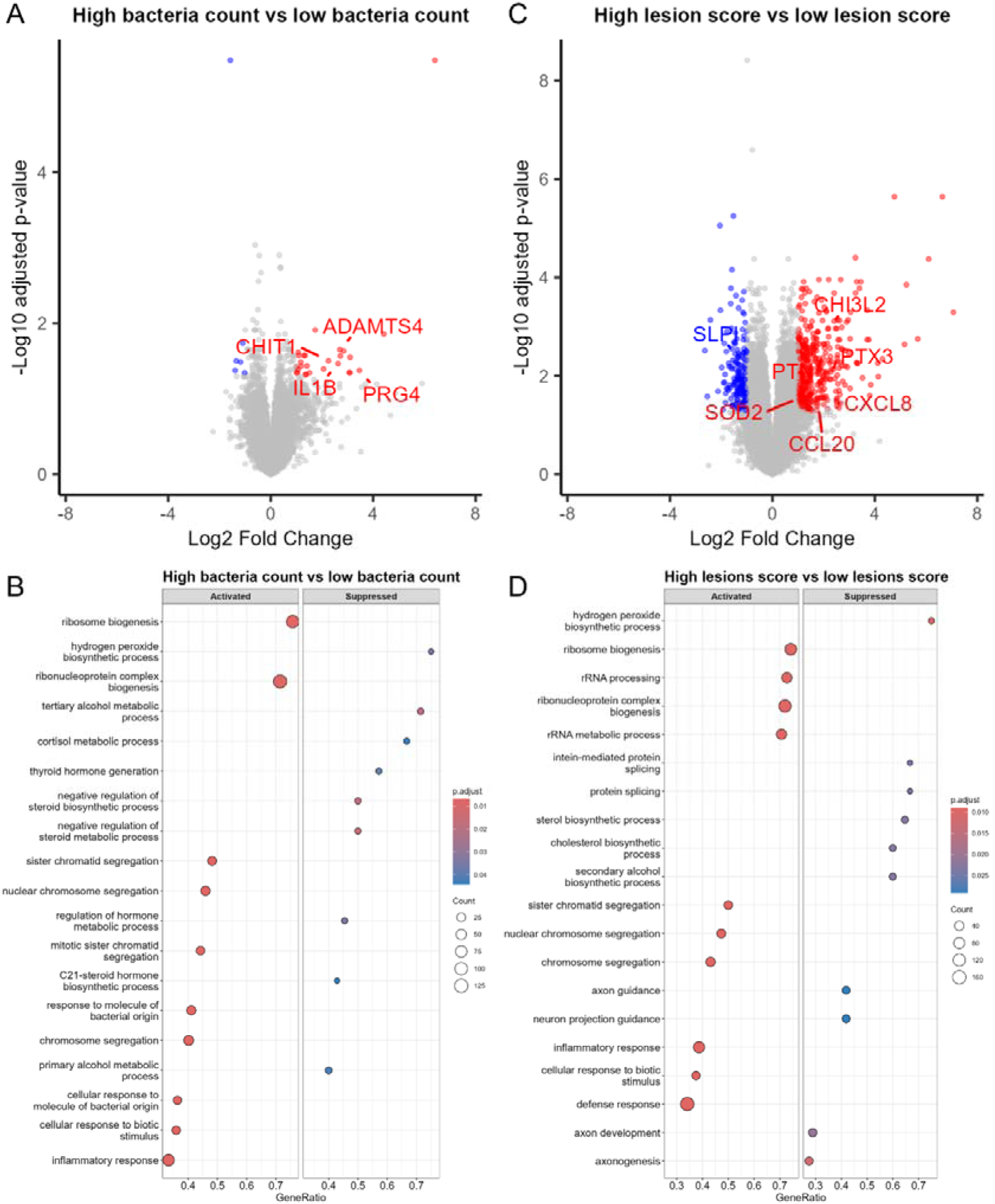
The effect of FLAMOD nebulization in the transcriptional landscape is transitory and changes in the transcriptome three days after *App* infection is driven by the severity of the lung infection. Pigs were infected with 1 x 10^7^ CFU *App* serotype 2 and treated daily with 4 mg/kg of penicilling G (*App*+PenG; n=9) or with one dose of 1.3 mg of FLAMOD in addition to the antibiotic therapy (*App*+PenG+FLAMOD; n=10). (A) Volcano plot showing differential gene expression between animals with a high bacterial count (log10 CFU>4) vs animal with a low bacterial count (log10 CFU <4). (B) Gene set enrichment analysis for differentially expressed genes between animals with high vs low bacterial counts. (C) Volcano plot showing differential gene expression between animals with a high lesion score (>6) vs animals with a low lesion score (<6). Red color indicates padj <0.05 and log2 fold change higher than 1. Blue color indicates padj <0.05 and log2 fold change lower than −1. (D) Gene set enrichment analysis for differentially expressed genes between animals with high vs low lesions score. The gene ontology term Biological Process (BP) was used with a p value cutoff of 0.05 and the Benjamini-Hochberg (BH) method to adjust for multiple comparisons. Data are representative of 9-10 animals per experimental group.

## Discussion

In the present study we demonstrate for the first time that co-administration of an antibiotic with FLAMOD enhances the therapeutic efficacy against severe porcine respiratory disease caused by *App*. In addition to reducing bacterial loads in the lungs, the combination therapy with FLAMOD improves the overall health of the animals. This was evidenced by increased weight, as well as a reduction in both lung inflammation and lung tissue damage, highlighting the potential of FLAMOD to not only enhance antibiotic effectiveness but also promote better recovery outcomes in infected pigs.

The growing concern regarding the emergence of antibiotic resistance and the failure in reaching therapeutic doses, often encountered during the treatment of unhealthy or stressed pigs (3, 24–27). highlights the need for increasing the efficacy of antibiotic therapy. Our findings show that stimulating innate immune responses with the adjunct FLAMOD therapy can enhance antibiotic effectiveness in this model of pig App infection. This finding is supported by several studies using murine models of antibiotic resistant pneumococcal pneumonia, where flagellin/FLAMOD treatment re-sensitized pneumococcus to antibiotics that would otherwise fail to clear the infection (15–17).

Several studies using the murine model show that administration of flagellin prior to or at the moment of infection can protect against bacteria by inducing the expression of antimicrobial peptides and several cytokines that will lead to the recruitment of immune cells to the lungs (8, 9, 28). We have observed a similar response in the pig, where flagellin was able to increase *CXCL8* expression, recruit immune cells to the lungs and improve the pig’s lungs defense against *Pseudomonas aeruginosa* (7). However, we did not observe a positive effect of FLAMOD prophylactic intervention in pig *App* infection in the first study. On the contrary, pigs pre-treated with FLAMOD 24 h pre-infection with *App* had a tendency towards a higher disease severity, with more affected lung tissue and more lungs showing typical pneumonia lesions in that group. This finding was surprising, as previous studies in murine models of pneumococcal infection revealed that prophylactic flagellin administration 12 to 24 h pre-bacterial challenge was protective to the animals (8), thus the timing of administration should have been optimal for protection. One could speculate that the increased immune response prior to infection could enhance the release of *App* virulence factors and endotoxins, however, this still remains to be investigated.

The protective effect of flagellin used post-infection in a therapeutic context have been previously described. Recent studies using a mouse model demonstrated that a combination of mucosal administered flagellin and a low-dose antibiotic, given either orally or intraperitoneally, was more effective against *Streptococcus pneumoniae* lung infections than standalone antibiotics (14–17). This approach reduced lung bacterial loads and mortality rates. It also proved effective in treating pneumococcal superinfection following influenza (14). Furthermore, flagellin had a synergistic effect on antibiotic therapy and was effective against amoxicillin-resistant *S. pneumoniae* (17). Our results show that the administration of flagellin post-infection in combination to a subtherapeutic dose of antibiotic significantly enhances protection in a pig *App* infection model. It is noteworthy that nebulization of just a single dose of flagellin into the lungs (in addition to antibiotics) resulted in a major decrease in the bacterial load, reducing it by 489-fold, compared to antibiotic therapy alone. This was accompanied by a diminished amount of lung lesions and a tendency to higher weight gain. Together these findings highlight the potential of flagellin as an adjunct to traditional antibiotic therapy in livestock.

The protective effect of flagellin is largely attributed to its capacity to initiate a transient inflammatory response. In the lungs, flagellin predominantly targets epithelial cells (5, 29), which express high levels of TLR5 (9). Studies in both mice [8, 10, 27] and pigs (7, 30) show that upon stimulation, epithelial cells can secrete bactericidal compounds, with a direct effect on the bacterial pathogen, as well as pro-inflammatory cytokines. The resulting proinflammatory environment leads to the recruitment and activation of immune cells, including neutrophils (7, 10) and dendritic cells (28). Neutrophils play a key role in the clearance of bacteria from the lungs (7, 28). Although one of the potential risks of the use of TLR agonists is the possibility of triggering an exacerbated and sustained inflammatory response, administration of moderated doses of flagellin have been reported to be safe and have not lead to such undesired effects (31, 32). The transcriptomic data at 3 days post-treatment support that FLAMOD does not increase inflammation and expression of inflammatory markers as cytokine/chemokine expression was not differentially modulated among the treatment groups. Previous data suggest that the local effect of flagellin in the lungs is transitory and prominent during the first 24h post-treatment in various animal models, including mice and pig (7, 15). In the present study, samples were collected 3 days post-infection, when the peak effect of flagellin on gene expression in the lungs has been surpassed. The notable variability in the pig lungs transcriptional response to *App* infection correlated to the severity of lesions. Indeed, animals presenting severe lesions had increased lung expression of inflammatory genes. This observation correlated with only 30% of the pigs treated with flagellin developing severe lesions compared to 66% in the PenG-treated group. Aerosolized FLAMOD also triggered a systemic effect as evidenced by an increase in the production of blood circulating cytokines. Such systemic stimulation of immune mediators has been previously described in mice treated with flagellin and could reveal the increased capacity of immune cell mobilization and maturation from the blood into the lungs (9, 33).

In conclusion, boosting innate immunity via flagellin administration can potentiate the therapeutic efficacy of antibiotics against *App* infections. This synergistic effect holds promise as a new therapeutic approach for controlling *App* infections in pigs, particularly in the context of the growing concerns surrounding the emergence of antibiotic resistant bacteria.

## Data availability

RNA-seq datasets have been deposited in GEO under the accesion number GSE253855

## Acknowledgements

The authors express their gratitude to the staff at the Unité Expérimentale de Physiologie Animale de l’Orfrasière INRAE–Research Centre of Tours (France) and the staff of the INRAE-PFIE animal experimental platform (UE-1277 PFIE, INRAE Centre de Recherche Val de Loire, Nouzilly, France) for their technical support.

## Author contributions

**IF**: Data curation, Formal analysis, Investigation; **VG**: Investigation, Methodology; **CB**: Investigation, Methodology, Resources; **MR**: Investigation, Methodology, Resources; **AD**: Investigation, Methodology, Resources; **AP**: Investigation, Methodology, Resources; **TB**: Data curation, Formal analysis, Supervision, Writing – review & editing; **MB**: Conceptualization, Investigation, Supervision, Writing – review & editing; **ST**: Resources, Funding acquisition, Writing – review & editing; **NS**: Conceptualization, Resources, Funding acquisition, Writing – review & editing; **JCS**: Conceptualization, Project administration, Resources, Writing – review & editing, Funding acquisition; **ICP**: Conceptualization, Data curation, Formal analysis, Investigation, Project administration, Supervision, Validation, Visualization, Writing - original draft-, Writing – review & editing, Funding acquisition.

