## supplementary for "Flagellin aerosol administration improves the efficacy of antibiotic treatment in *Actinobacillus pleuropneumoniae* infected pigs"

**Supplementary material and methods**

**Flow cytometry**

Broncho-alveolar lavages (BAL) were obtained post-mortem by instilling and retrieving two 15-mL aliquots of sterile high glucose DMEM containing 100 IU/ml penicillin, 2.5 µg/ml amphotericin and 50 µg/ml gentamicin. Following collection, red blood cells were lysed using an erythrocyte lysis buffer and washed with DMEM supplemented with 10% FBS and 100 IU/ml penicillin before staining for cell surface markers.

Cell population in the BAL were characterized by their specific surface markers as described in (22). Cell surface staining was performed in PBS-EDTA supplemented with 5% porcine serum and 5% goat serum for 30 min in ice. Primary antibodies were used as follows: anti-MHCII (mouse IgG2a, clone MSA3, dilution 1/250; Kingfisher Biotech); anti-CD172a (mouse IgG2b, clone 74-22-15a, dilution 1/500; BD Biosciences); anti-CD163 PE-conjugated (mouse IgG1, clone 2A10/11, dilution 1/20; BIO-RAD). Secondary antibodies anti-mouse IgG2a coupled to PE-Cyanine7 and anti-mouse IgG2b couple to APC-Cyanine 7 were purchased from Invitrogen and used at a 1/200 dilution. Cell viability was assessed using the Fixable Viability Dye eFluor™ 450 (ThermoFisher). Between labellings cells were washed with PBS-EDTA. BAL samples were acquired on a BD LSR Fortessa X-20 (BD Biosciences, San Jose, CA, USA) and data was analyzed using Kaluza analysis software v2.1. Neutrophils and alveolar macrophages were classified as described previously (22) where neutrophils were defined as MHCII^null^/CD163^null^/CD172a^+^, while alveolar macrophages were defined as MHCII^+^/CD163^+^/CD172a^+^.

**Real-time quantitative PCR (RT-qPCR)**

Gene expression was evaluated from the cranial and caudal left lung lobes. Briefly, RNA was purified from using the NucleoSpin^®^ RNA XS kit (Macherey-Nagel, Düren, Germany). The extracted total RNA was converted to cDNA by the reverse transcription of 20 ng of RNA with iSCRIPTtm Reverse Transcription Supermix (Bio-Rad). RNA quality and integrity were assessed by calculating A_260_/A_280_ in NanoDrop spectrophotometer analysis (NanoDrop Technologies, Wilmington, USA) and capillary electrophoresis (Agilent 2100 Bioanalyzer, Agilent Technologies, Inc., Santa-Clara, USA). Pre-amplification was performed with cDNA diluted 1:10 in TE (10 mM Tris, 0.1 mM EDTA, pH 8.0, Fluka Biochemika #93283) with Preamp Master Mix (Fluidigm, # 100-5581). The pre-amplification thermal cycle conditions were: 95 °C for 2 min followed by 14 cycles of 95 °C for 15 seconds and 60 °C for 4 min. Pre-amplified cDNAs were diluted 1:5 in TE (Tris-EDTA buffer solution BioUltra, for molecular biology, pH 8.0, 10 mM Tris-HCl, 1 mM disodium EDTA Sigma-Aldrich #93283) after a treatment with Exonuclease I (20u/µl, New England Biolabs) for 30 min at 37°C followed by 15 min at 80°C. Real-time quantitative PCR was performed in a 48 x 48 Dynamic Array Integrated Fluidic Circuit (Fluidigm) with the primers described in Table S1, 2x SsoFast EvaGreen with Low Rox (BioRad 172-5211) and 20 X DNA binding Dye (Fluidigm PN 100-7609). The thermal cycle conditions were 1 min at 95°C, followed by 30 cycles of denaturation for 5 s at 96°C and annealing/elongation for 20 s at 60°C. Data were analyzed using the 2^ΔΔCt^ method and the results are expressed as relative fold change.

**Supplementary table 1. Primers used for evaluation of real-time quantitative PCR.**

| Gene | Forward primer | Reverse primer |
| --- | --- | --- |
| B2M | CGAGACCACTAACCGGCATCA | TGGATTCATCCAACCCAGATGCA |
| RPL19 | GCTCAGAAGATACCGTGAATCTA | ACACATTCCCCTTCACTTTCA |
| CXCL8 | TGCCTTCTTGGCAGTTTTCC | CTGCACTTACTCTTGCCAGAAC |
| PTN | GCAACTCCAAAATGCAGACCC | ACACACACTCCACTGCCATT |
| PTX3 | GTTGGCCGAGAACTCCGAT | GAAGCATGCTCTCCCGCATC |
| SLPI | CAAGTGCACAAGTGACTGGC | GGCCATAGACCACTGGACAC |
| SOD2 | GGCCTACGTGAACAACCTGA | TGATTGATGTGGCCTCCACC |
| TNFAIP3 | AGGAAGCTTGTGGCACTGAA | TCCTGAACACCCCACATGTAC |

Annealing temperature is 60°C for all primer pairs.

**Supplementary Figure 1. Flow cytometry evaluation of the percentage of neutrophils and alveolar macrophages in the BAL.** Neutrophils were identified as MHCII^null^CD172a^+^CD163^null^ while of alveolar macrophages in the BAL identified as MHCII^+^CD172a^+^CD163^+^. A pairwise Wilcoxon test with Holm correction for multiple comparisons was used to evaluate significant differences between the FLAMOD+*App*, *App* and *App*+PenG groups*.* Triangle shapes indicate animals that presented lung lesions. Data are representative of 24 animals.

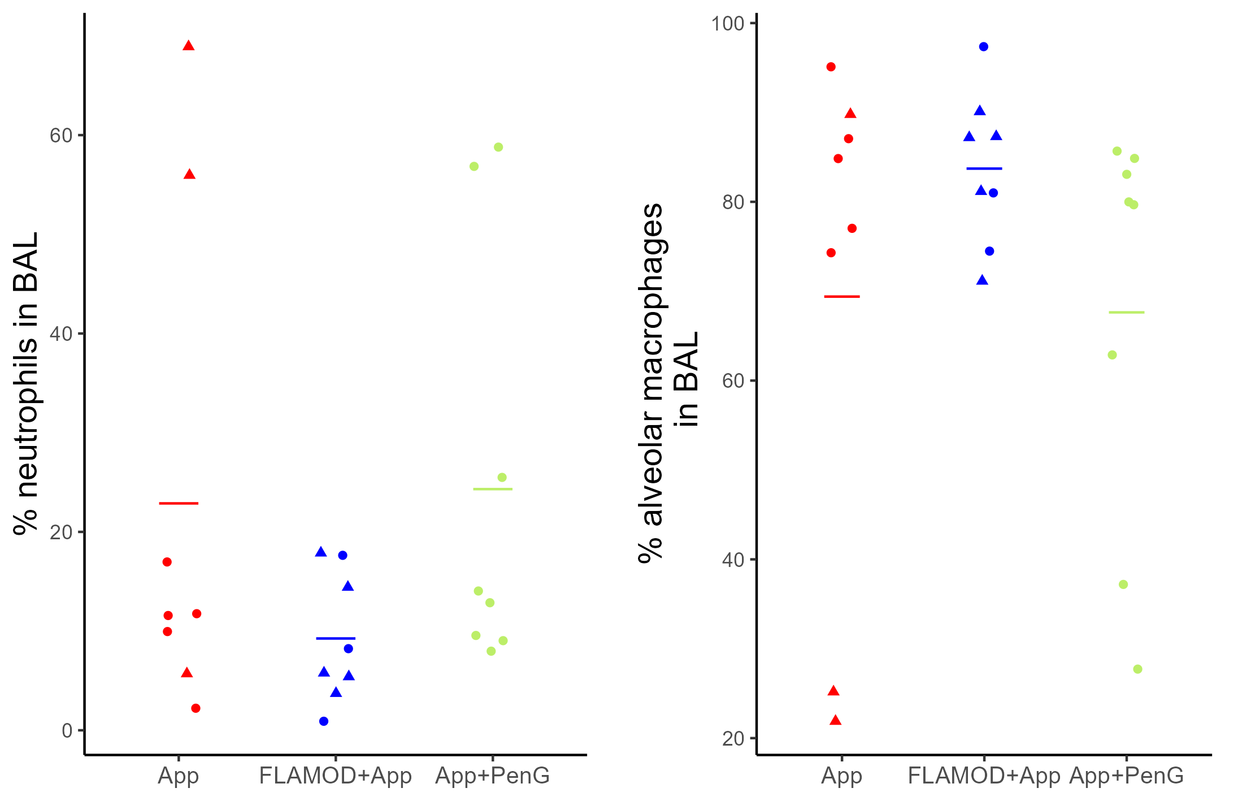

**Supplementary Figure 2. Effect of the combination therapy of FLAMOD and penicillin G on the pig circulating cytokines during *App* infection.** One-month old pigs were infected with 1 x 10^7^ CFU *App* serotype 2 and treated daily with 4 mg/kg of penicilling G (*App*+PenG; n=9) or with one dose of 1.3 mg of FLAMOD in addition to the antibiotic therapy (*App*+PenG+FLAMOD; n=10). Measurement of IL-1RA and TNF production in the serum at 0, 6 and 72h p.i. A mixed-effects linear model was applied to analyze statistical differences among the treatment groups, temporal factors, and the interaction effect in the time series data. Pig identification was included as a random effect in the model.

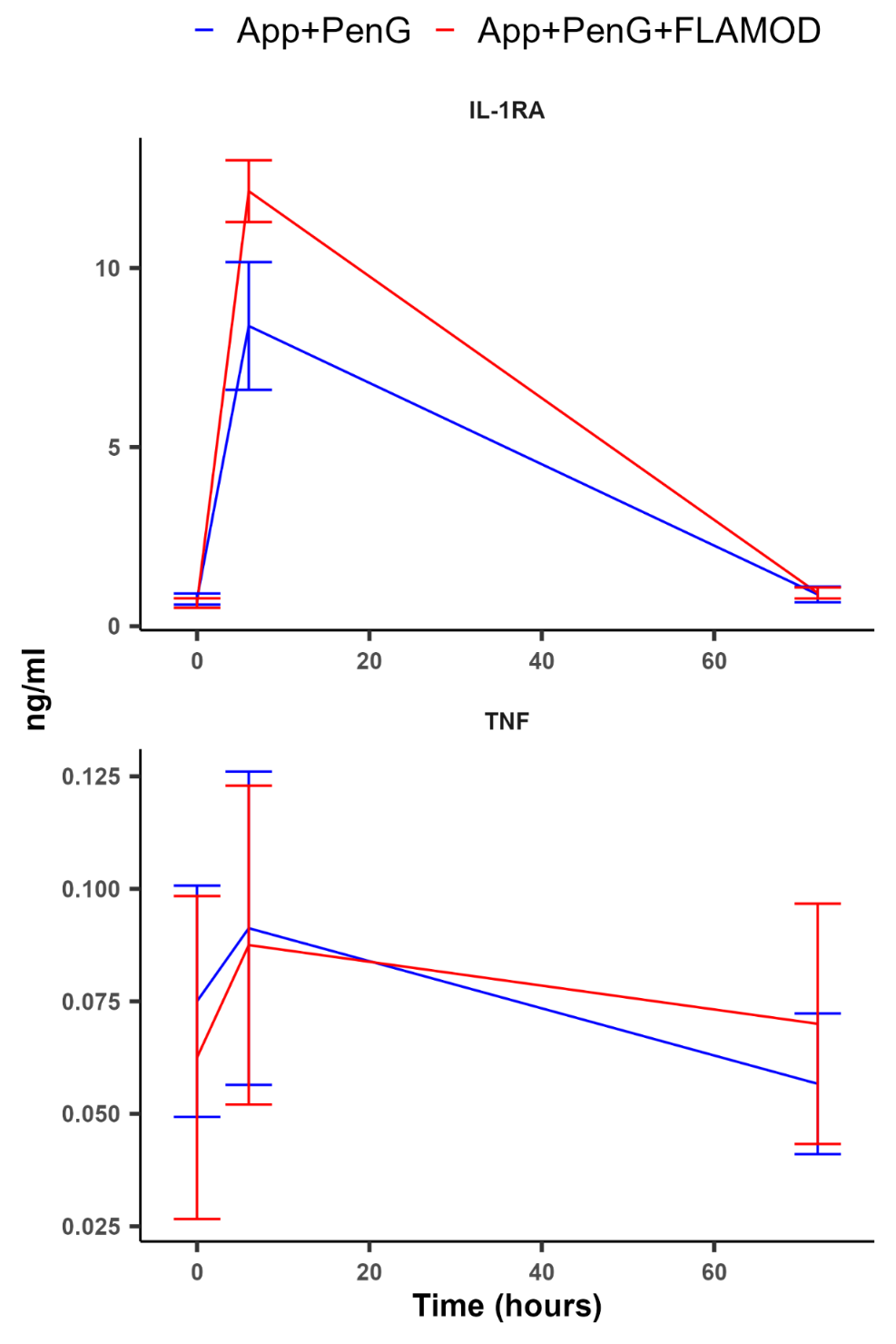

**Supplementary Figure 3. Effect of FLAMOD nebulization in the transcriptional landscape of infected lungs.** Volcano plot showing differential gene expression between animals treated with *App*+PenG vs *App*+PenG+FLAMOD. Data are representative of 9-10 animals per experimental group

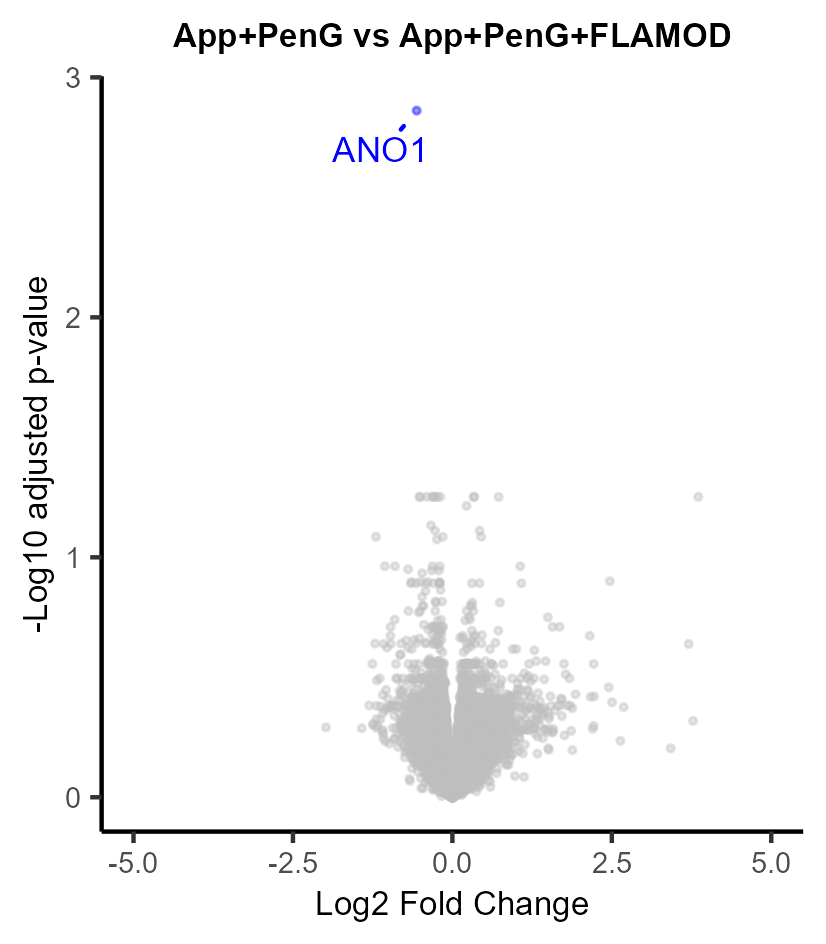

**Supplementary Figure 4. Effect of FLAMOD nebulization in the transcriptional landscape of infected lungs.** Quantitative PCR analysis from the cranial and caudal left lobes of pigs presenting high and low lesion score. Gene expression is shown relative to the low lesion score samples for each lung lobe. Differences were evaluated by using the Mann-Whitney Wilcoxon test. Data are representative of 9-10 animals per experimental group.

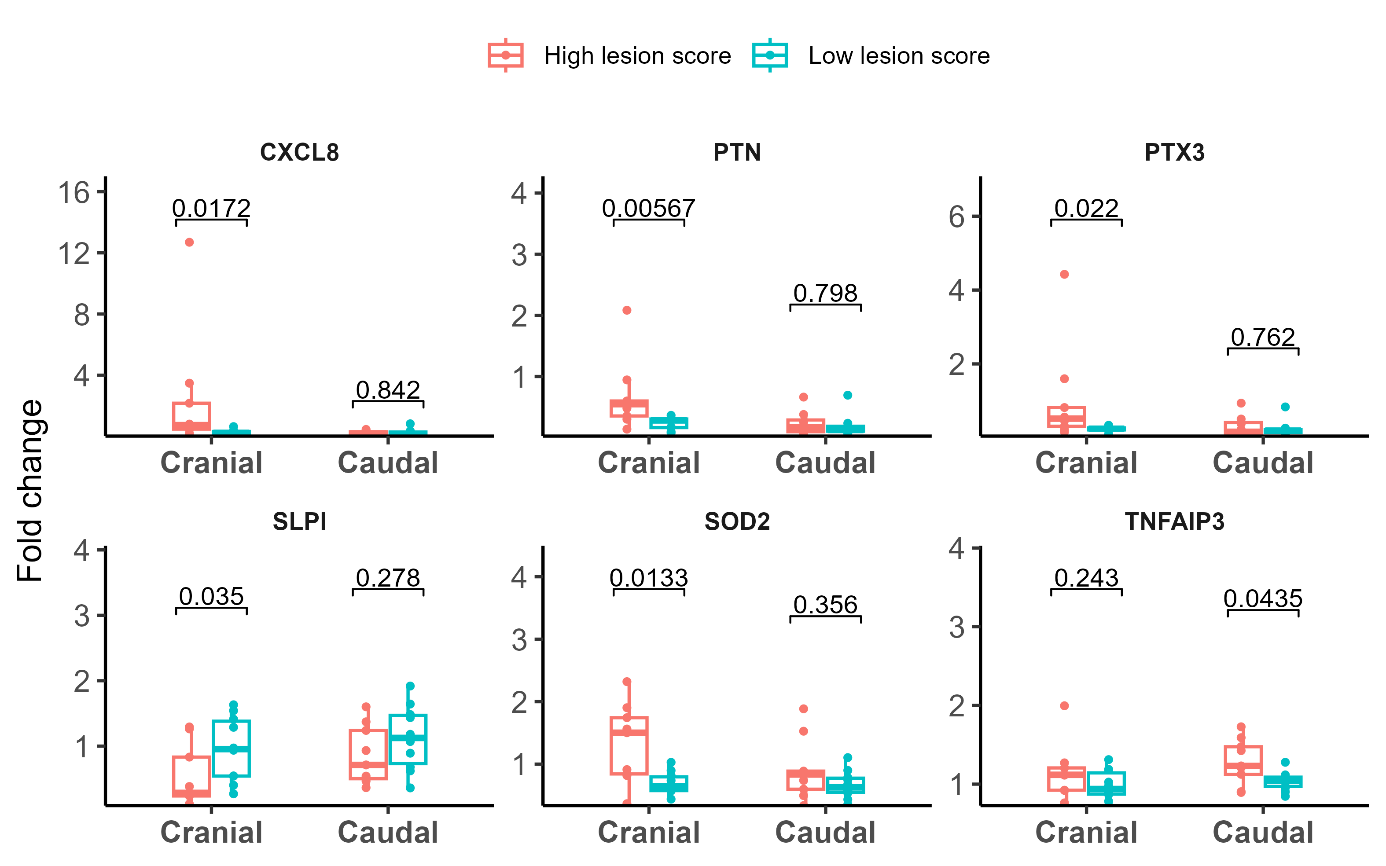
